# Consensus Label Propagation with Graph Convolutional Networks for Single-Cell RNA Sequencing Cell Type Annotation

**DOI:** 10.1101/2022.11.23.517739

**Authors:** Daniel P Lewinsohn, Katinka A Vigh-Conrad, Donald F Conrad, Cory B Scott

## Abstract

**Motivation:** Single-cell RNA sequencing (scRNA-seq) data, annotated by cell type, is useful in a variety of downstream biological applications, such as profiling gene expression at the single-cell level. However, manually assigning these annotations with known marker genes is both time-consuming and subjective.

**Results:** We present a Graph Convolutional Network (GCN) based approach to automate the annotation process. Our process builds upon existing labeling approaches, using state-of-the-art tools to find cells with highly confident label assignments through consensus and spreading these confident labels with a semi-supervised GCN. Using simulated data and two scRNA-seq data sets from different tissues, we show that our method improves accuracy over a simple consensus algorithm and the average of the underlying tools. We also compare our method to a non-parametric neighbor majority approach, showing comparable results. We then demonstrate that our GCN method allows for feature interpretation, identifying important genes for cell type classification. We present our completed pipeline, written in PyTorch, as an end-to-end tool for automating and interpreting the classification of scRNA-seq data.

**Availability:** Our code for conducting the experiments in this paper and using our model is available at https://github.com/lewinsohndp/scSHARP

**Supplementary information:** Supplementary data are available at *Journal Name* online.

## Introduction

Single-cell RNA sequencing (scRNA-seq) measures the amount of RNA from each gene present in an individual cell, serving as a proxy for gene expression. High-quality labels of cell type (based on the transcriptional profile produced by scRNA-seq) have proven valuable for characterizing gene expression of cells, and for discovering cell types and genetic drivers of disease. Traditionally, these labels are produced by unsupervised clustering followed by labeling clusters with known marker genes. However, unsupervised clustering is limited by issues such as the size of scRNA-seq data sets as well as subjectivity in reclustering and biological interpretation of clusters [1].

The limitations of traditional cell type annotation methods have necessitated the development of automated methods for cell labeling. Three main categories of tools have emerged: marker gene-based, correlation-based, and supervised classification-based [2]. Marker-based approaches employ known marker genes for labeling, while correlation and supervised learning-based approaches require manually labeled scRNA-seq data sets with the cell types of interest. Within these broad categories, the performance of individual tools varies widely across data sets [3]. As a result, using the consensus of multiple classification tools could yield higher accuracy. However, there currently exist no tools for researchers to easily apply multiple classification algorithms to their scRNA-seq data. Even once different predictions have been collected, deciding how to handle cells with ambiguous predictions between tools remains an open question.

We address these issues in two ways. First, we provide a pipeline for annotation of scRNA-seq data with multiple state-of-the-art annotation algorithms. Second, we implement and test a semi-supervised Graph Convolutional Network (GCN) as a mechanism to propagate labels from confidently labeled cells to unconfidently labeled cells. We compare our method to five state-of-the-art cell classification tools, along with a simple max consensus and a non-parametric neighbor majority approach. We show our method improves overall classification accuracy (and specifically, classification accuracy on unconfidently labeled cells) compared to component tools and the simple consensus of the tools. We also demonstrate our approach shows comparable performance with the non-parametric neighbor majority approach.

Following accurate classification, an important aspect of model usage within the biological sciences is interpretability. There has been significant development around neural network interpretation, resulting in a variety of approaches for assigning attribution scores to input features. Gradient-based attribution methods for neural networks have proven valuable given their efficiency and ease of implementation [4]. Importantly, trained neural network models can be queried for important features for classification that are not otherwise obvious. One effective algorithm for gradient-based interpretation is DeepLIFT. This method decomposes the output prediction of a neural network on a specific feature by backpropagating through the network. DeepLIFT tries to explain to the difference between the output and a neutral reference output, generated by a neutral reference input [5].

We demonstrate the use of DeepLIFT [5] as an effective interpretation tool for our GCN model, allowing researchers insight into classification decisions and important cell type gene markers. The ability to apply DeepLIFT to our model is a valuable feature compared to other baseline methods used in this study. Specifically, we show interpretation of our trained model identifies differentially expressed genes with biological significance not explicitly used as markers. Furthermore, we show adding these highly-attributed genes as markers for marker-based tools can improve accuracy.

## Methods

### Our Model

#### Selected Component Methods

Our method relies on picking confident labels for a subset of cells in a data set. To do this, our pipeline currently includes five different state-of-the-art annotation methods: SCINA [6], ScType [7], scSorter [8], SingleR [9], and scPred [10]. We briefly describe each of these tools and refer the reader to these tools’ original papers for more details. ScType employs a clustering-based approach that inputs the scRNA-seq gene expression matrix along with cell type gene markers, and outputs predictions [7]. scSorter is another clustering-based approach that uses the scRNA-seq gene expression matrix along with cell type gene markers. This method recognizes that over-expression of certain marker genes is not present in populations of many cell types and attempts to address this problem [8]. SCINA is another state-of-the-art approach that uses the same information as ScType and scSorter, with the added benefit of being much faster. SCINA uses an expectation-maximization algorithm to assign labels [6]. SingleR requires both the scRNA-seq gene expression matrix and a labeled reference gene expression matrix. This method uses correlation between the reference and query sets to extend labels [9]. scPred requires the same information as SingleR. This method uses feature reduction to pull out important cell type features and then a probabilistic machine learning algorithm for label prediction [10]. We note that our pipeline could be applied to a broader set of tools.

#### Picking Confident Labels via Consensus

We briefly describe our pipeline for generating “confident” i.e. consensus cell labels. Gene counts are initially filtered to first remove cells expressed in fewer than 200 genes and then filtered to remove genes expressed in fewer than 3 cells. Tools make predictions on further preprocessed gene counts as specified by the given tool, returning an *n* by *t* matrix *R* where *t* is the number of annotation tools and *Ri,j* is the prediction from tool *j* for cell *i*. We then one-hot encode the predictions from each column of *R* resulting in *t* matrices of size *n* by the number of cell types. We then add these matrices to construct a new matrix *E* where the rows correspond to cells and columns correspond to the amount of votes for each cell type. Finally, we create confident labels *c ∈* ℝ^*n*^ where *cn* corresponds to the predicted cell type for cell *n* if a majority of tools agreed and -1 if not. Selecting confident labels entails setting a threshold of confidence which we set at .51 for a simple majority. For cell *xi* we find *mi* = *max*(*Ei/sum*(*Ei*)) and assign label -1 if *mi <* .51 and the label of the corresponding column of *mi* if *mi >*= .51. This results in *c ∈* ℝ^*n*^ containing the confidently labeled and unconfidently labeled cells.

#### Semi-supervised GCN

Our data is presented to our model as a preprocessed scRNA-seq matrix for *n* cells, denoted by *X* = *{x*_1_, …, *x*_*n*_} *⊆* ℝ^*F*^ where *F* is the number of features for each cell. We use Principal Components Analysis (PCA) to project the cell-by-gene matrix *X* into a 500D vector space. To train our GCN, we draw random minibatches of size b at a time. We construct *G* as the k-nearest neighbor (k-NN) graph of the cells included in each minibatch with *k* neighbors, including self-loops. We experimented with different values of *k*, finding lower neighbors and batches performed better. We input *G* into the following GCN architecture.

EdgeConv [11] is a graph convolution operator developed for prediction from point clouds that captures local structure in graphs while remaining permutation invariant. We construct a GCN with *l* EdgeConv layers with SiLU [12] activation function and summation aggregation. The SiLU activation function is a common neural network activation function that we found to work well for our GCN architecture. A final linear layer projects node embeddings into label space (whose dimension is the number of cell types in our data set). For architecture details see Appendix A. We train our GCN for 150 epochs, with the Adam optimizer [13] at an initial learning rate of 0.0001. Our training loss is Cross-Entropy loss on the set of confidently labeled cells. Nearest neighbors are generated separately for each batch of each epoch. We consider multiple choices for each of the hyperparameters batch size *b*, neighbors *k*, number of message passing steps *l*, and final embedding layer size *e*, selecting the hyperparameters that result in best performance on held-out validation data. See Experiment Settings for details.

#### Interpretation with DeepLIFT

We employ DeepLIFT [5] with the Rescale rule as implemented by Captum [14], a model interpretability and understanding library for PyTorch. The Rescale rule is one option for assigning the contribution scores of the immediate inputs to each neuron. We use the same hyperparameters batch size *b*, neighbors *k*, number of message passing steps *l*, and final embedding layer size *e* as used during training. Please refer to the Experiment Settings section for how these hyperparameters are chosen. DeepLIFT requires a reference input to determine the reference activation of a neuron. We use a reference count matrix of all zeros and the same size as our scRNA-seq count matrix *X*. We then use our tool’s predictions for each cell type as its target neuron for calculating the gradient. This results in attribution matrix *A* with the same size as *X*. We use *A* to calculate the mean feature attribution scores for each gene, for all cells of each type. We then take the absolute value of these attributions to compute a general importance score per feature/gene. Note that our transformed DeepLIFT scores do not indicate negative or positive importance for each feature, just overall importance. Finally, we scale each cell type to have unit variance to allow comparison of gene importance scores across cell type. Calculating these attribution scores is only possible for a differentiable model, like our GCN – not possible for any of the underlying tools. We note here that the pipeline we analyzed with DeepLIFT included the PCA step, resulting in attribution scores per gene for each cell.

### Non-parametric Neighbor Majority Approach

In addition to the max consensus baseline described below in Experiment Settings, we implemented a non-parametric neighbor majority approach. This is similar to the message passing step in our GCN model, with the difference that this method does not use a neural network to encode/decode messages. This method operates on the 500D vectors produced as the principal components of the preprocessed gene expression matrices for each data set. We first construct a k nearest neighbors graph in Euclidean distance, using a batch size of 1000. During each round of message passing, the unconfidently labeled nodes’ labels are updated as the majority label of their k nearest neighbors in euclidean distance (only considering those neighbors who have been labeled thus far). We test three strategies during hyperparameter optimization for convergence of this method:

- one round of label propagation;
- iterating until less than 5 percent of labels change between epochs; and
- iterating until all cells are labeled or 50 epochs have gone by.

We show accuracy scores across a variety of k values and convergence strategies in Appendix D.

### Data Sets

#### Preprocessing Data

The initial input to our pipeline is a scRNA-seq count matrix *X* where *Xng* corresponds to observed gene counts for gene *g* in cell *n*. Cells expressing *<* 200 genes and genes expressed in *<* 3 cells are removed, and *X* is row-normalized according to *x*_*i*_,: = *log*(1 + ((*x*_*i*_,: *∗* 10000)*/* Σ _*j*_ *x*_*ij*_)). Both of these steps are common scRNA-seq preprocessing steps [15][16]. Finally, we use Principal Components Analysis (PCA) to project *X* down to 500 features per cell.

#### Prototyping Data

Separate simulated data that was generated with a different seed than the test simulated data was used for developing our model. We also used a subset of the PBMC data that was not included in the final test set to develop our model. We approached this task as a transductive semi-supervised graph learning task where each scRNA-seq data set is its own graph. However, no ground truth labels are provided for the learning task. We did not run our model on our four test data sets until architecture details were finalized. Additionally, only confidently labeled nodes are used for hyperparameter optimization in these test sets, meaning no ground truth labels are involved. It is also important to note no trained model weights carried over from the prototyping data sets to the testing data sets.

#### Simulated Data

We generated our simulated data sets with Splatter [17] and parameters estimated from 4000 Pan T Cells from a healthy donor [18]. Each simulated data set contains 1000 cells, evenly split between four cell types with different transcriptomic profiles. To demonstrate that our pipeline can also incorporate reference-based tools (like SingleR and scPred), we also generated 1000 reference cells for each data set with the same gene profiles (separated from the original data with a batch.facScale of 0.5 to simulate batch effects). Simulated data sets vary by the de.facScale parameter which determines the magnitude of differential expression in differentially expressed genes between cell type groups. Five markers were selected randomly from the top ten differentially expressed genes from each cell type. The 0.7 de.facScale and 0.8 de.facScale simulated data sets had 156 and 85 unconfidently labeled cells respectively.

#### Real Data

It is not usually feasible to acquire ground truth labels for scRNA-seq data. An alternative gold standard is Fluorescence-activated Cell Sorting (FACS), which pre-sorts cells by markers prior to conduction of scRNA-seq [19]. We test our model on two FACS-labeled data sets.

First, we use a scRNA-seq data set generated from mouse testis cells [20]. This data contains three cell types: 292 Spermatogonia, 244 Spermatocytes, and 156 Spermatids after filtering. Cell type markers were selected from relevant literature [21] and a full list of markers is included in Appendix I. Additionally, no reference data set was used for this data. There were 46 unconfidently labeled cells after prediction tool voting on the data set.

Second, we use an scRNA-seq data set generated from Human peripheral blood mononuclear cells (PBMCs) [22]. This data contains ten cell types, however, we removed cell types not purely sorted by FACS, combined CD4+ T cells, and combined CD8+ T Cells. This resulted in five cell types: 9,106 B Cells, 2,341 Monocytes, 7,572 Natural killer (NK) Cells, 38,006 CD4+ T Cells, and 19,856 CD8+ T Cells. We used the same markers as scSorter [8] with the addition of CD8A, CD8B, and CD4 to distinguish CD4 and CD8 T Cells. A full list of markers is included in Appendix I. We used the 10X PBMC 3k data set as a reference as in [23]. This data set contained 2,310 unconfidently labeled cells.

## Results

### Accuracy on Test Sets

#### Experiment Settings

For the simulated and testis data sets, 20 percent of confidently labeled cells were masked and held out as a validation set. We performed a hyperparameter optimization search (see Appendix A for details) for options of batch size *b*, neighbors *k*, layers *l*, and embedding layer size *e*, selecting the GCN architecture with the highest validation accuracy. For the PBMC data set, five percent of the data set was used for this same hyperparameter optimization process as using the entire data set was computationally intractable. Each GCN used EdgeConv feature propagation between each node and its *k* closest neighbors, with distance determined dynamically between node features (including the PCA features at the first layer). Five random initializations of the optimal model were then trained for 150 epochs as described above, and mean accuracy was recorded. We use max consensus, the non-parametric neighbor majority approach, and other tool accuracies as baselines for our model. Max consensus simply chooses the cell type with the most votes. In the event of a tie, this method returns “unknown”. The k hyperparameter and convergence method for the non-parametric neighbor majority approach was chosen with the same validation set used for GCN hyperparameter optimization. We report total accuracy and unconfident cell accuracy for each data set. “Unconfident cell accuracy” refers to the accuracy on only the cells where the underlying tools did not find consensus.

#### Simulated Data Sets

For both simulated data sets, the optimal model has batch size 20, 2 nearest neighbors, and 2 EdgeConv layers. For the simulation with 0.7 de.facScale, 25-dimensional embedding space was optimal, whereas for the simulation with 0.8 de.facScale the optimal value was 40. Additionally, 15.6% and 8.5% of cells were unconfidently labeled in the 0.7 de.facScale and 0.8 de.facScale data sets respectively. Table 4 shows our model outperforms all other methods for total accuracy and slightly under performs SingleR for unconfident cell accuracy on the 0.7 de.facScale data.

The non-parametric neighbor majority approach proves extremely ineffective in propagating confident labels to unconfidently labeled cells in both simulated data sets. This could be due to the transcriptional similarity of all four cell types in the simulated data sets. We compare the average distance between each cell type cluster in Appendix H, finding all cell types in each simulated data set are very transcriptionally similar. In contrast, the testis and PBMC data sets show varying levels of transcriptional dissimilarity depending on which cell types are compared. This suggests that in data sets with many similar cell types, the non-parametric neighbor majority approach would fail.

#### Testis Data Set

Only marker-based prediction tools were used for this data set as no labeled reference was easily available, and we wanted to test our method without reference-based tools. The optimal model for this data set used batch size 20, 2 nearest neighbors, 2 EdgeConv layers, and embedding layer size of 25. Importantly, 6.6% of cells were unconfidently labeled by the component tools. Table 4 shows accuracy results, demonstrating our GCN model outperforms all other methods for both total and unconfident cell accuracy, except for the non-parametric approach. The accuracy of the non-parametric approach is within the standard deviation of our model for both the total and unconfident accuracies.

These results suggest both our GCN model and the non-parametric neighbor majority effectively propagate consensus labels to unconfidently labeled cells in this testis scRNA-seq data. This is clear given the improved accuracy over all component methods and significant improvement of our model over the tool average accuracy by 14.1%. Our model therefore demonstrates improved accuracy over component methods and comparable accuracy to the non-parametric neighbor majority approach, with the major advantage of being interpretable.

#### PBMC Data Set

The optimal model for this data set used batch size 50, 2 nearest neighbors, 2 EdgeConv layers, and embedding layer size of 25. 3.0% of cells were unconfidently labeled by the component tools. Table 4 shows accuracy results. For accuracy on unconfident cells, the GCN model places third behind ScType and the non-parametric approach. For overall accuracy, our model ties with the non-parametric approach to the third significant digit. Additionally, the accuracy of the non-parametric approach is well within the standard deviation of our model accuracy on the unconfident set.

These accuracy results highlight the ability of our GCN model and the non-parametric neighbor majority approach to propagate confident labels in the PBMC data. We again show a clear improvement over all component methods, demonstrating the value of this approach in labeling scRNA-seq cells.

### Feature Interpretation

Figure 2A shows the five most highly attributed genes by DeepLIFT for each cell type in the testis data set. Interestingly, all of these top genes have uniquely high attribution in their important cell type. The highly attributed genes for a cell type also have relatively high gene expression in that cell type as indicated by Figure 2B. We also observe high expression of Spermatocyte genes in Spermatid cells. It is important to note DeepLIFT interpretation indicates importance of both input marker genes and genes not explicitly given to marker-based tools. For example, DeepLIFT indicates genes Ncl, Ldhc, and Tnp1 that are differentially expressed in those cell types, but not explicitly included as marker genes. Tsga8 and Acrv1 are both Spermatid marker genes that appear in the top ten highest attributed DeepLIFT genes. Additionally, Spermatogonia marker gene, Dazl, is both a used marker gene and highly attributed DeepLIFT gene. A full list of marker genes used and a list of the top ten highest attributed DeepLIFT genes for each cell type are included in Appendix I and Appendix G, respectively.

**Fig. 1.**
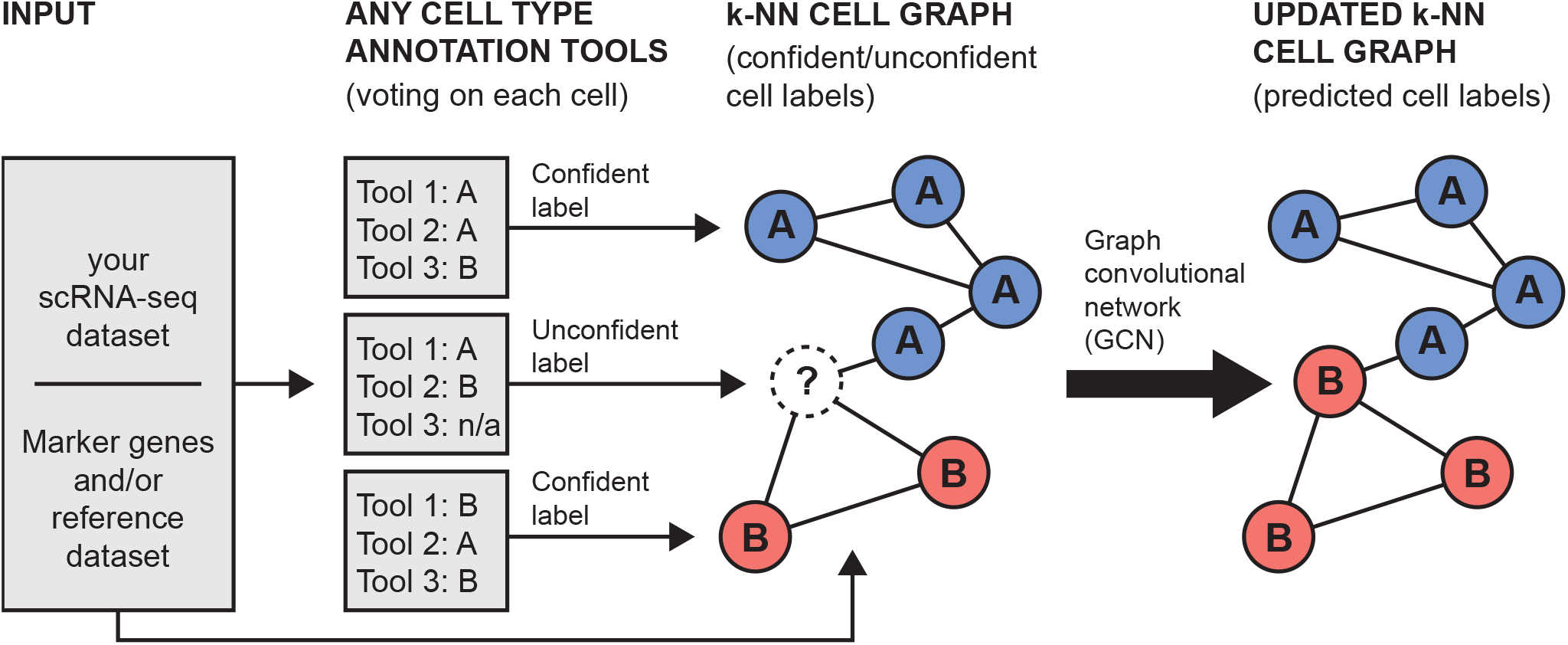
Starting from an scRNA-seq data set, a user can apply any number of tools which classify cell types (for example, classification from a pre-labeled reference data set or from reference genetic markers). If a majority of these underlying tools assign the same label to a cell, we say the ensemble is confident in this label. Our GCN learns to propagate labels from confidently labeled cells to the rest of the cells (“unconfident cells”) via message passing in a K-nearest neighbor graph.

**Fig. 2.**
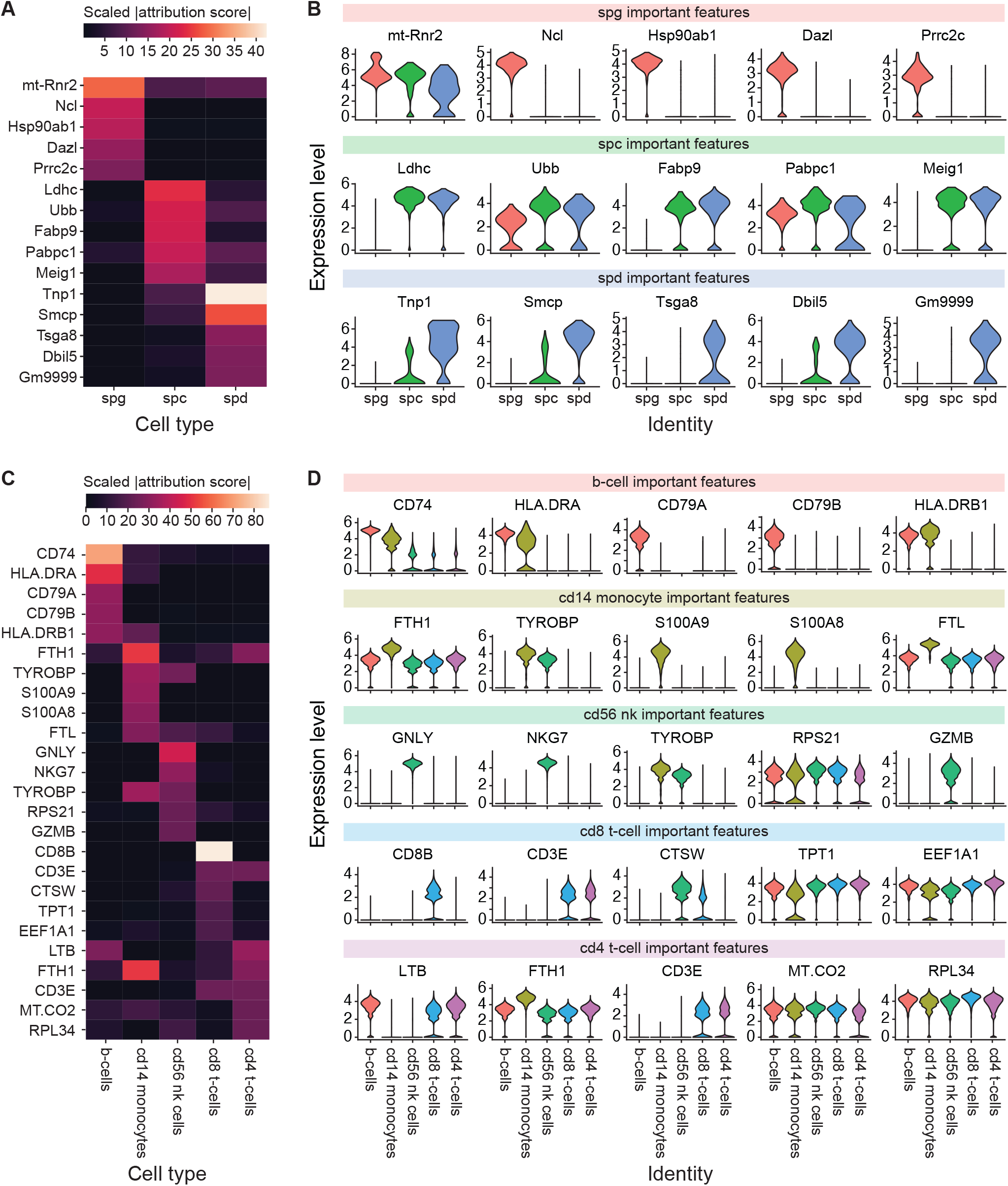
Heatmap of DeepLIFT attribution scores after absolute value and scaling by cell type for top five most important features by predicted cell type and violin plot of log normalized expression for each gene by predicted cell type. a. Attribution heatmap for testis data set. b. Expression plots for testis data set by predicted cell type. c. Attribution heatmap for PBMC data set. d. Expression plots for PBMC data by predicted cell type.

Figure 2C shows the DeepLIFT attribution scores for the top five highly-attributed genes by cell type for the PBMC data. Figure 2D shows actual gene expression of these highly attributed genes in the data set. For B Cells, Monocytes, and NK Cells we see a clear connection between the genes picked out as important by DeepLIFT and the genes expressed by those cell types. However, for CD4 and CD8 T Cells, the expression is not clearly higher for all genes. Importantly, we do observe CD8B as the most important gene for CD8 T Cell classification, a key marker for the cell type. We also observe CD3E (another important marker for all T Cells) as an important gene for both sub types of T Cells. However, we do not observe CD4 as highly attributed in CD4 T Cells. This result is not totally unexpected given that CD4 is only measured by scRNA-seq in a relatively low percent of CD4 T Cells [24]. Additionally, the DeepLIFT scores for CD4 and CD8 T Cells may be less informative because the GCN often misclassifies CD8 T Cells as CD4 T Cells. This is likely due to the large heterogeneity and general transcriptional similarity of T Cell subtypes. Finally, DeepLIFT again reveals both used marker genes and novel genes. Specifically, DeepLIFT highly attributes CD74, FTH1, GNLY, and more genes that are differentially expressed, but not originally included as marker genes. DeepLIFT also highly attributes original marker genes such as CD79A, CD8B, CD3E, and CD3D. Again, a full list of PBMC marker genes is included in Appendix I.

We also compared highly attributed DeepLIFT genes in both data sets to genes identified in differential expression analysis. This experiment and the top ten highest ranked genes for each method can be found in Appendix G. In both data sets, both DeepLIFT and differential expression analysis revealed many common genes. However, there are also many genes not jointly identified by both methods. We suggest the use of DeepLIFT as a complimentary analysis to differential expression analysis, providing another method for identifying candidate cell type marker genes.

Overall, DeepLIFT interpretation proves effective in both the testis and PBMC data sets, revealing novel genes not originally included as markers. Importantly, the GCN is the only one of our tested methods that can be interpreted using DeepLIFT. This adds a valuable feature for researchers to both further understand classification decisions and get insight into novel cell type markers.

### Highly Attributed Genes Improve Marker-Based Methods

We added the top three highly attributed genes from DeepLIFT for each cell type not already included as a marker gene to the list of marker genes. We then re-ran the marker-based tools ScType, scSorter, and SCINA with these markers on the testis and PBMC data sets to compare tool accuracy. This experiment shows the value of highly attributed DeepLIFT genes in assisting future classification.

Table 2 shows the change in accuracy before and after adding highly attributed genes in the testis data set. Both scSorter and SCINA show significant improvement in accuracy with the addition of highly attributed DeepLIFT genes. scSorter shows large in improvement in Spermatocytes while SCINA shows extreme improvement in Spermatids, demonstrating the effectiveness of using DeepLIFT attributed genes across different cell types and tools. ScType accuracy remains the same across all cell types. Although ScType does not improve, it is important to note accuracy does not go down with the addition of these new marker genes.

**Table 1.** Accuracy scores (percentage of cells correctly classified) for all data sets for both all cells and “unconfident cells”: cells for which the underlying methods did not have consensus. GCN accuracies are the mean *±* standard deviation of accuracies from five randomly initialized trials. Simulation 0.7 and Simulation 0.8 are referred to as 0.7 de.facScale and 0.8 de.facScale data sets respectively.

| Method | Simulation 0.7 |  | Simulation 0.8 |  | Testis |  | PBMC |  |
| --- | --- | --- | --- | --- | --- | --- | --- | --- |
|  | All | Unconf. (156) | All | Unconf. (85) | All | Unconf. (46) | All | Unconf. (2,310) |
| Ours (GCN) | **90.0 ± .61** | 66.0 ± 3.9 | **96.1 ± .22** | **83.3 ± 2.6** | 86.2 ± .12 | 80.9 ± 1.8 | **93.1 ± .05** | 70.6 ± 1.5 |
| Max Consensus | 86.4 | 42.9 | 91.2 | 25.9 | 80.8 | 0.0 | 91.2 | 6.8 |
| Tool Avg. | 69.3 ± 14 | 33.1 ± 32 | 76.7 ± 12 | 34.6 ± 28 | 72.1 ± 16 | 21.7 ± 36 | 75.0 ± 6.1 | 36.9 ± 34 |
| ScType | 64.8 | 13.5 | 79.9 | 29.4 | 84.8 | 63.0 | 85.5 | **81.8** |
| scSorter | 85.8 | 51.3 | 88.7 | 38.8 | 77.5 | 2.2 | 71.5 | 65.9 |
| SCINA | 53.7 | 7.7 | 58.7 | 2.4 | 53.9 | 0.0 | 73.3 | 6.2 |
| SingleR | 83.1 | **80.1** | 85.2 | 77.6 | *NA* | *NA* | 70.3 | 13.4 |
| scPred | 59.2 | 12.8 | 71.0 | 24.7 | *NA* | *NA* | 74.4 | 17.1 |
| Non-Parametric | 82.3 | 16.7 | 90.3 | 15.3 | **86.3** | **82.6** | **93.1** | 70.8 |

**Table 2.** Change between accuracy score before and after adding highly attributed genes as markers in testis data set. Accuracy difference is recorded for each cell type: Spermatogonia (SPG), Spermatocytes (SPC), and Spermatids (SPD).

| Method | SPG | SPC | SPD |
| --- | --- | --- | --- |
| ScType | 0.00 | 0.00 | 0.00 |
| scSorter | 4.11 | 12.71 | 2.56 |
| SCINA | 7.53 | 2.87 | 33.97 |

Table 3 shows the change in accuracy before and after adding highly attributed genes in the PBMC data set. ScType accuracy remains the same for B Cells, NK Cells, CD4 T Cells, and CD8 T Cells. For Monocytes, ScType accuracy slightly decreases. scSorter accuracy increases in B Cells, Monocytes, NK Cells, and CD4 T Cells. In CD8 T Cells, accuracy decreases. SCINA accuracy increases in all cell types except CD4 T Cells, showing large increases in Monocytes, NK Cells, and CD8 T Cells. In general, we observe increased or maintained improvement in B Cells, Monocytes, and NK Cells, indicating the effective use of DeepLIFT attributed genes as markers in these cell types. However, both T Cell subtypes show varying performance, with decreases in accuracy in both scSorter and SCINA. We believe this is due to the inaccuracy of our model in distinguishing these T Cell subtypes as previously discussed. This suggests using DeepLIFT attributed genes falls short in cell types with poor model accuracy. We suggest further validation of all genes with direct expression of genes in the scRNA-seq data to confirm new markers.

**Table 3.**
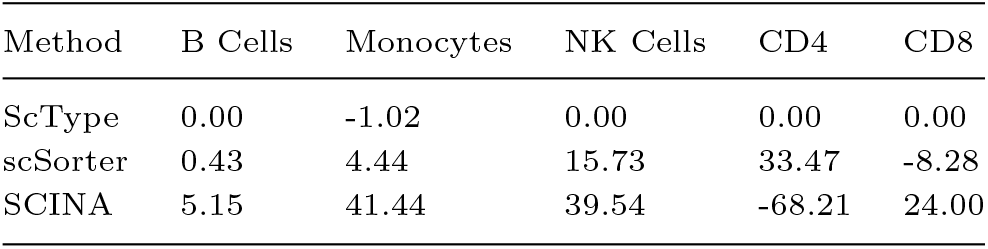
Change between accuracy score before and after adding highly attributed genes as markers in PBMC data set. Accuracy difference is recorded for the following cell types: B Cells, Monocytes, Natural Killer Cells (NK Cells), CD4 T Cells (CD4), and CD8 T Cells (CD8)

| Method | B Cells | Monocytes | NK Cells | CD4 | CD8 |
| --- | --- | --- | --- | --- | --- |
| ScType | 0.00 | -1.02 | 0.00 | 0.00 | 0.00 |
| scSorter | 0.43 | 4.44 | 15.73 | 33.47 | -8.28 |
| SCINA | 5.15 | 41.44 | 39.54 | -68.21 | 24.00 |

## Discussion

scRNA-seq cell type annotation remains an open and essential problem in bioinformatics. Accurate cell type labels are critical for downstream analysis and robust hypothesis testing with single-cell data. In this work we propose a novel framework for scRNA-seq cell type annotation that improves upon existing approaches. Building upon existing annotation tools, we implement an EdgeConv based GCN model to propagate consensus based confident labels to the remaining unlabeled cells. We show an improvement in accuracy over a baseline max consensus algorithm and the average tool accuracy. We also show similar, if not improved, performance over a non-parametric neighbor majority propagation approach. We then demonstrate the ability to identify important genes for classification via model interpretation with DeepLIFT. The model interpretation is especially valuable for researchers as it has the potential to uncover novel gene markers and provide insight into the model’s decisions.

We provide evidence for the accuracy of our GCN approach through comparison to state-of-the-art methods ScType, scSorter, SCINA, SingleR, and scPred on a variety of data sets, both simulated and real, including both Mouse and Human tissues. Our method improves accuracy in all data sets over all of these methods, providing strong evidence for the value of our method. A key takeaway from our research is the value of using a consensus of multiple annotation tools to improve general classification accuracy in scRNA-seq data.

We further provide evidence for the interpretability of our trained model with DeepLIFT. Highly attributed DeepLIFT genes include both given markers and novel genes that are differentially expressed in the actual data. This interpretation method is only possible for our method out of all compared methods, and provides a valuable tool for researchers when analyzing their scRNA-seq data.

One interesting aspect of our research is the similarity in classification accuracy of the non-parametric neighbor majority approach and our GCN approach for both real data sets. This suggests for these two data sets, the majority of valuable information for classification is simply in the existing labels of each cell’s neighbors. However, the GCN approach does improve performance when multiple cell types are transcriptionally similar (as in the simulated data sets). This suggests the value of using our GCN approach emerges when transcriptional profiles are similar across multiple cell types. These findings are relevant to both this project and the general function of message passing in graphs. As a whole, we find the most value in our GCN approach given its consistency in classification accuracy across all tested data sets and interpretability.

Overall, our method will serve as a valuable tool for researchers. They will be able to easily apply five state-of-the-art labeling tools and obtain accurate labels with our GCN propagation. Importantly, our pipeline accommodates any set of predictions, allowing our tool to be applied to any cell type with or without a reference data set.

## Acknowledgments

This work was supported by the National Institutes of Health [P50 HD096723-01]. Thanks to the Colorado College Department of Mathematics and Computer Science for supporting this research via the faculty summer research stipend. The authors thank Eisa Mahyari for valuable conversations. The authors thank William Holtz, Benjamin Modlin, and Max Perozek for their hard work on the scSHARP and R4scSHARP software packages.

## Data Availability

All used real scRNA-seq data sets are publicly available and cited above. Our code for conducting the experiments in this paper and using our model is available at https://github.com/lewinsohndp/scSHARP.

## A Hyperparameter Search Details

We performed a hyperparameter search over the number of neighbors *k*, batch size *b*, number of GCN layers *l*, and final embedding layer size *e*. Specifically, we tested number of neighbors at 2, 5, and 10, batch size at 20, 35, 50, 65, 80, and 95, GCN layers from 2 to 4, and final embedding sizes of 15, 25, 40, and 60. We tested GCN models on a validation set, consisting of 20 percent of the confidently labeled cells, for each combination of these hyperparameters. Figure 3 shows the validation accuracy distributions at each hyperparameter in each data set. Generally, we see high validation accuracy with lower neighbors and two or three GCN layers. Batch size and final embedding size seem to have a wide distribution of validation accuracy. For each data set, the combination of hyperparameters with the highest validation accuracy was selected for testing.

**Fig. 3.**
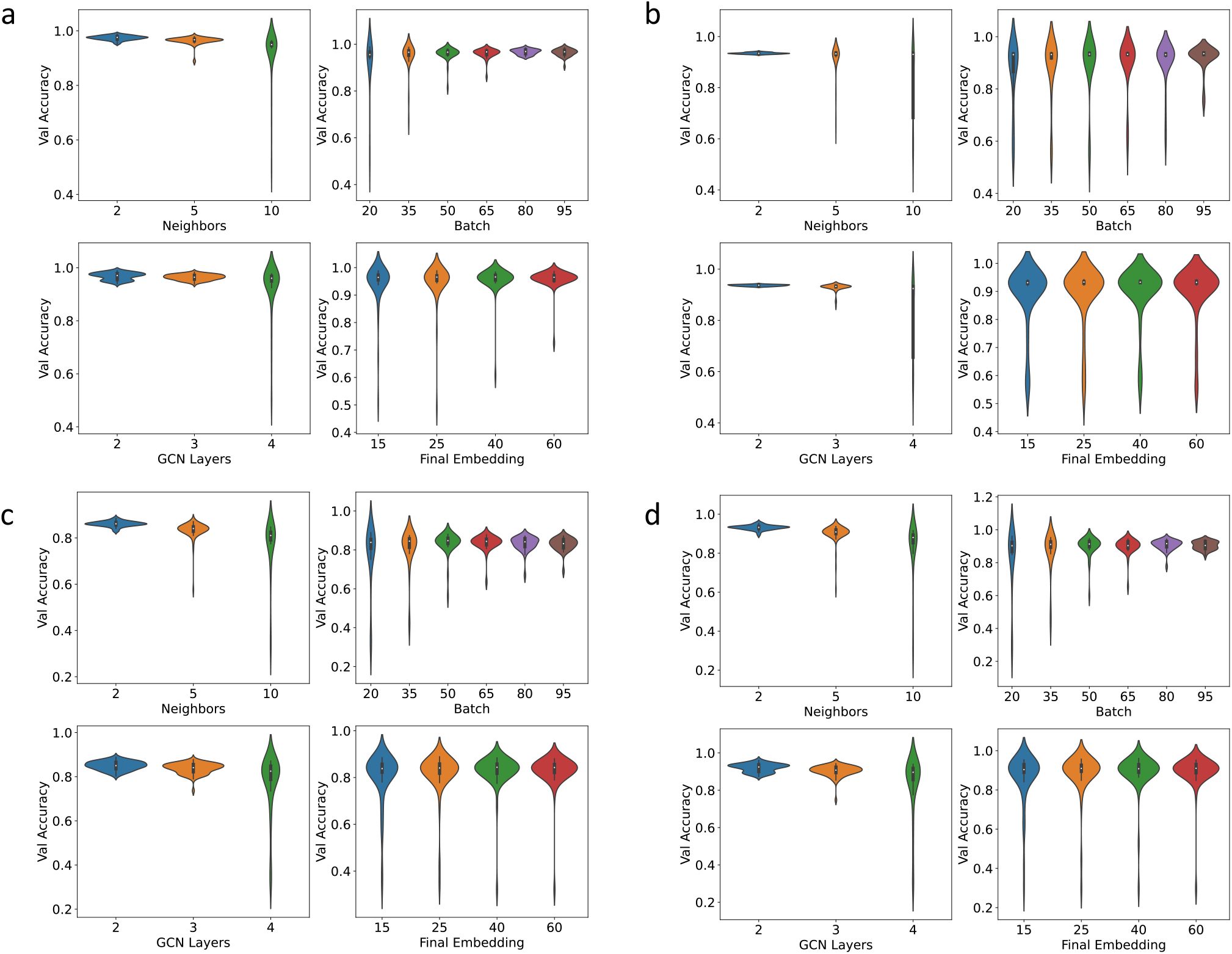
Spread of validation accuracy scores as a function of various hyperparameters. The hyperparameters included are number of neighbors, batch size, GCN layers, and final embedding layer size. a. Testis data set. b. PBMC data set. c. Simulation 0.7 data set. d. Simulation 0.8 data set de.facScale Value

Our model architecture consists of *l* EdgeConv layers. Each EdgeConv layer consists of one round of message passing along edges of the graph, followed by a dense neural network model that maps from one layer’s embedding space to the next layer’s. Each node aggregates information using the sum of its received messages (from neighbors and itself). In all of our model architectures, the first layer takes input embedding size 500 and outputs embedding size 1000. The middle layers accept embedding size 1000 and output embeddings of the same size. The final layer accepts embedding size 1000 and outputs final embedding size *e*. Both hyperparameters number of layers *l* and final embedding size *e* are included in the hyperparameter search.

## B de.facScale Simulation Parameter

See Figure 4 for details on how the de.facScale parameter affects classification difficulty. With de.facScale *≤* 0.5, the generated data is too difficult for any of the component tools to analyze – all of the component tools do poorly. On the other hand, de.facScale *≥* 0.9 is too easy – the cell types are well-separated enough in gene space that all component methods are able to classify them correctly. We generated simulated data with de.facScale values of 0.7 and 0.8, as these values produce a challenging, but still attainable, benchmark for classification.

**Fig. 4.**
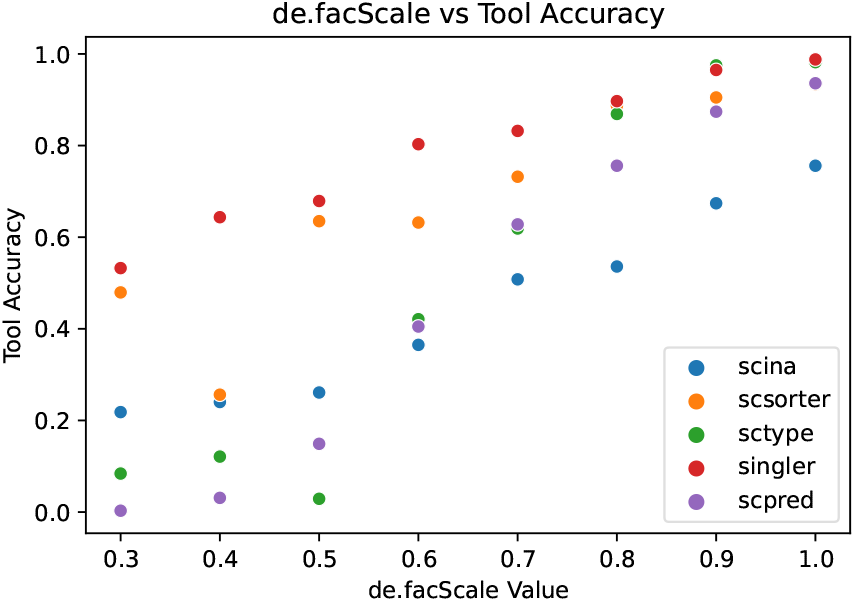
Plot showing relationship between de.facScale simulation parameter and component tool accuracy. ScType, SCINA, ScSorter, SingleR, and ScPred were included as component tools.

## C GCN Confusion Matrices

See Figure 5 for confusion matrices from GCN predictions on all data sets. We note that nearly all of these confusion matrices are characterized by a single cell type being the majority of unconfident cells. It is unclear why this is the case for the PBMC and testis data. In the synthetic data, one possible explanation is the way we choose marker genes. We randomly select five of the top ten differentially expressed genes in the simulated data of each cell type as markers (this is standard practice for using Splatter). It is possible this method results in some cell types with better markers than others. This was intentional, as in real-world data not all cell types will always have the same strength of cell type marker. For the actual data sets, this is likely because certain cell types (such as CD4 and CD8 T Cells) are more transcriptionally similar and likely to be misclassified. A common theme in both of these cases is that within each data set, some cell types are inherently easier to classify than others.

**Fig. 5.**
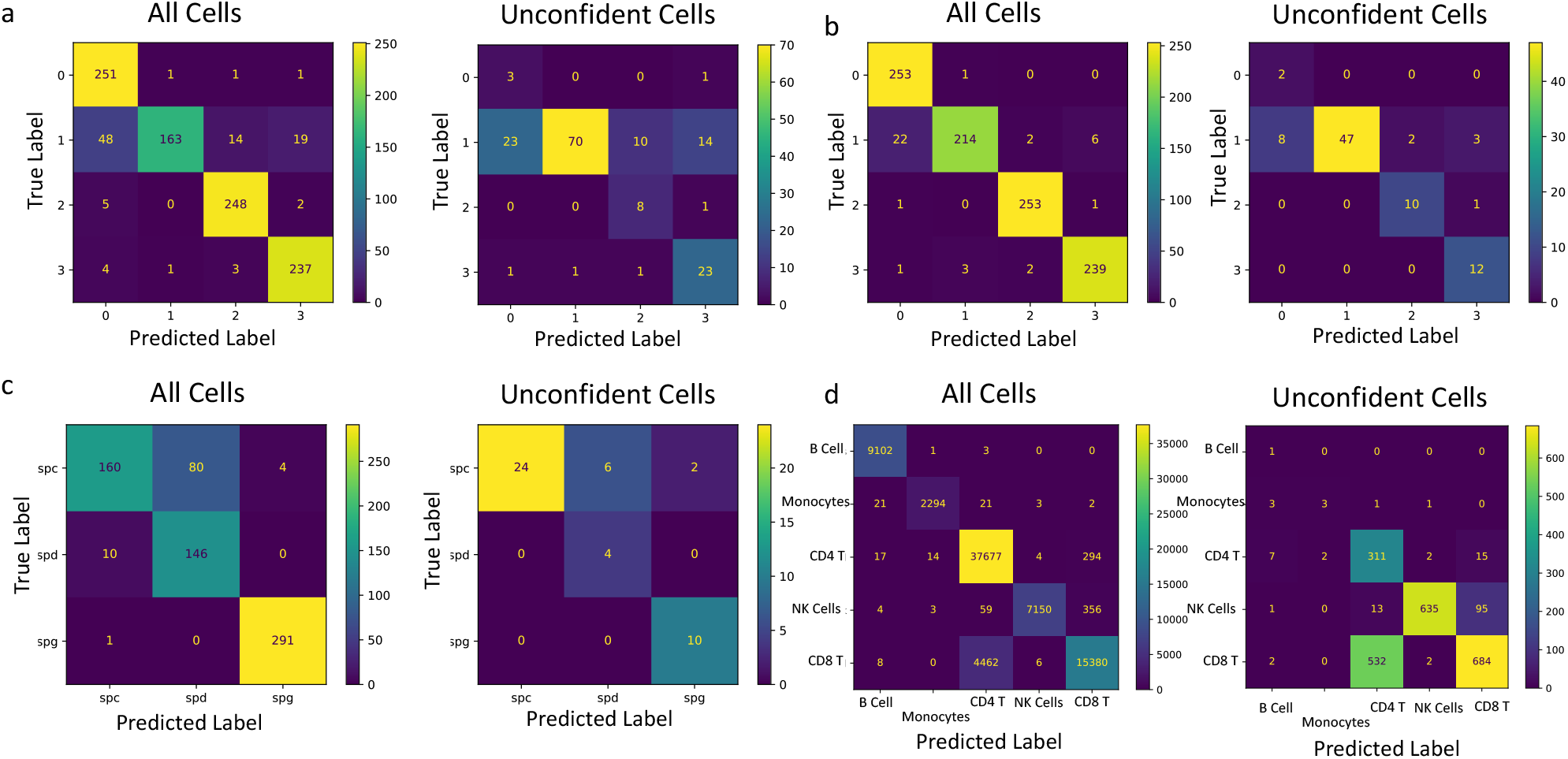
Confusion matrices for GCN predictions in all and unconfidently labeled cells. a. Simulation 0.7 b. Simluation 0.8 c. Testis d. PBMC

## D Non-parametric Neighbor Majority Label Propagation

The results for the neighbor-majority label propagation approach across a variety of k values are shown in Figure 6. For each data set, we see that the convergence method does not greatly impact the accuracy for a given neighbor hyperparameter. We do see the neighbor hyperparameter is essential for the accuracy of this method.

**Fig. 6.**
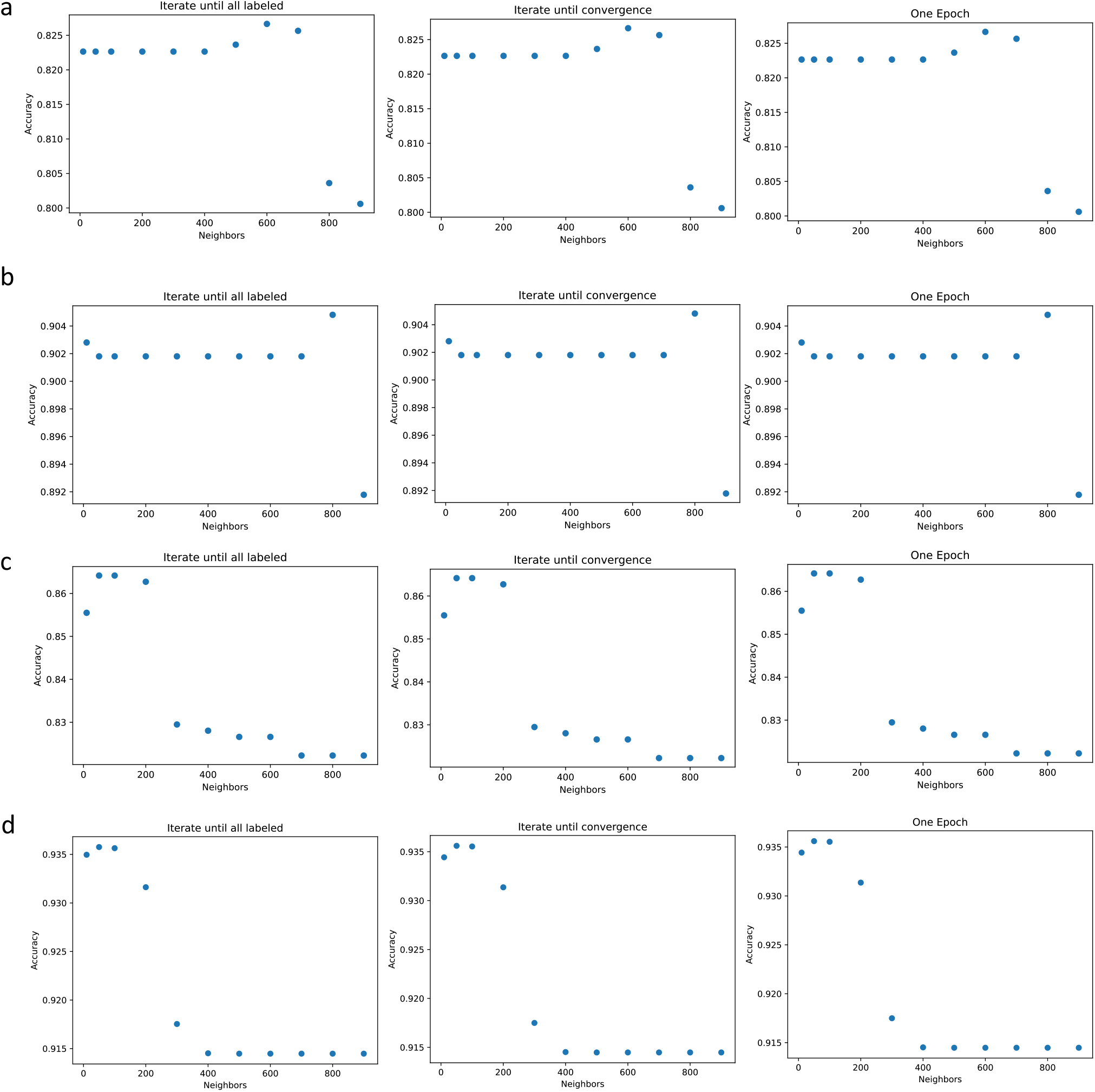
Number for neighbors vs accuracy score for different convergence methods and data sets. a. Simulation 0.7 b. Simulation 0.8 c. Testis d. PBMC

## E Testing in Simulated Data with Class Imbalance

To further test how our model performs in data sets with class imbalance, we generated two additional simulated data sets. These data sets were generated in the same way as the de.facScale 0.7 and 0.8 data sets, except proportions of 0.1, 0.2, 0.3, and 0.4 were used to generate the number of cells for each cell type. This resulted in 374, 318, 204, and 104 cells in Groups 1, 2, 3, and 4, respectively, in both data sets.

We then ran the same experiments as described in Experiment Settings and reported results in Table 4. Our model continues to out perform all underlying tools on overall accuracy in both data sets. Our GCN model also outperforms both the max consensus approach and the non-parametric neighbor majority approach.

**Table 4.** Accuracy scores (percentage of cells correctly classified) for simulated class imbalance data sets for both all cells and “unconfident cells”: cells for which the underlying methods did not have consensus. GCN accuracies are the mean *±* standard deviation of accuracies from five randomly initialized trials.

| Method | Simulation 0.7 Imbalance |  | Simulation 0.8 Imbalance |  |
| --- | --- | --- | --- | --- |
|  | All | Unconf. (70) | All | Unconf. (26) |
| Ours (GCN) | **94.0 ± .31** | 62.0 ± 4.4 | **99.5 ± .11** | **90.0 ± 4.4** |
| Max Consensus | 91.8 | 30.0 | 98.1 | 34.6 |
| Tool Avg. | 78.8 ± 9.9 | 34.9 ± 28 | 88.2 ± 10 | 38.5 ± 32 |
| ScType | 75.3 | 25.7 | 95.4 | 42.3 |
| scSorter | 87.1 | 62.9 | 95.4 | 76.9 |
| SCINA | 70.4 | 10.0 | 74.9 | 3.8 |
| SingleR | 91.4 | **65.7** | 95.5 | 61.5 |
| scPred | 70.0 | 10.0 | 79.8 | 7.7 |
| Non-Parametric | 92.1 | 34.3 | 98.1 | 34.6 |

These results in conjunction with the class imbalance seen in both the testis and PBMC data sets provide evidence for the robustness of our method in data sets with class imbalance.

## F Testing in Simulated Data with Variation in Number of Marker Genes

To explore how varying the number of marker genes for each cell type affects both the marker-based component tools and our model, we created two more simulated data sets. These data sets were generated in exactly the same way the de.facScale 0.7 and 0.8 data sets were generated. We then ran all component tools with a full set of marker genes (five for each cell type) and with a subset of these marker genes. In the reduced marker gene set, Group 1 had only two marker genes, Group 2 had three, Group 3 had four, and Group 4 had all five marker genes. The non-parametric approach was not included as it is not directly relevant to testing how varying the number of marker genes affects the component tools and our model. Additionally, the same model hyperparameters used for the original simulated data sets were used for this experiment.

Tables 6 and 7 show the by cell type change in accuracy for each of the marker-based tools as a result of reducing the marker gene set. For scSorter, we see a large decrease in accuracy for Group 1, where the most marker genes were removed, for both data sets. For SCINA, we see very similar results with both sets of marker genes, except for Group 2 in the Simulation 0.7 de.facScale data set where there is a large increase in accuracy for Group 2. ScType accuracies remained the same across both gene sets and both data sets. Table 5 shows the accuracy results for both our Simulated 0.7 and 0.8 de.facScale data sets with all and reduced marker genes. In both data sets with reduced marker genes, we see decreased max consensus accuracy, indicating the tool consensus is worse when the marker genes are reduced. As a result, we see decreased accuracy in our model as well when the marker gene set is reduced.

**Table 5.** Accuracy scores (percentage of cells correctly classified) for simulated marker variation data sets for both all cells and “unconfident cells”: cells for which the underlying methods did not have consensus. GCN accuracies are the mean *±* standard deviation of accuracies from five randomly initialized trials. Simulation 0.7 and 0.8 All refer to the data sets where all five markers were used. Simulation 0.7 and 0.8 Reduced refer to the data sets where markers were reduced as described in Appendix F.

| Method | Simulation 0.7 All |  | Simulation 0.7 Reduced |  | Simulation 0.8 All |  | Simulation 0.8 Reduced |  |
| --- | --- | --- | --- | --- | --- | --- | --- | --- |
|  | All | Unconf. (148) | All | Unconf. (193) | All | Unconf. (50) | All | Unconf. (87) |
| Ours (GCN) | **96.2 ± .23** | **89.5 ± 1.6** | **91.8 ± .51** | **80.2 ± 2.4** | **98.8 ± .20** | **94.0 ± 4.0** | **98.1 ± .11** | **92.9 ± 1.3** |
| Max Consensus | 89.8 | 45.9 | 82.0 | 29.0 | 96.0 | 38.0 | 93.0 | 34.5 |
| Tool Avg. | 64.7 ± 24 | 35.1 ± 24 | 63.7 ± 20 | 35.1 ± 25 | 75.7 ± 24 | 37.2 ± 25 | 73.3 ± 22 | 36.6 ± 21 |
| ScType | 66.5 | 38.5 | 66.5 | 43.5 | 85.7 | 36.0 | 85.7 | 47.1 |
| scSorter | 81.9 | 46.6 | 68.3 | 25.9 | 95.7 | 70.0 | 84.1 | 33.3 |
| SCINA | 25.6 | 2.0 | 33.9 | 3.6 | 34.9 | 0.0 | 34.8 | 1.1 |
| SingleR | 88.3 | 64.9 | 88.3 | 71.5 | 87.3 | 40.0 | 87.3 | 52.9 |
| scPred | 61.5 | 23.6 | 61.5 | 31.1 | 74.8 | 40.0 | 74.8 | 48.3 |

**Table 6.** Change between accuracy score before and after changing the number of markers used for each cell type in the Simulation 0.7 data sets. Accuracy difference is recorded for each cell type: Group 1 (1), Group 2 (2), Group 3 (3), and Group 4 (4). Group 1 had only two markers, Group 2 had only three markers, Group 3 had only four markers, and Group 4 had all five original marker genes.

| Method | 1 | 2 | 3 | 4 |
| --- | --- | --- | --- | --- |
| ScType | 0.00 | 0.00 | 0.00 | 0.00 |
| scSorter | -49.82 | -6.34 | -7.31 | 15.0 |
| SCINA | 0.00 | 36.65 | 0.39 | 0.42 |

**Table 7.** Change between accuracy score before and after changing the number of markers used for each cell type in the Simulation 0.8 data sets. Accuracy difference is recorded for each cell type: Group 1 (1), Group 2 (2), Group 3 (3), and Group 4 (4). Group 1 had only two markers, Group 2 had only three markers, Group 3 had only four markers, and Group 4 had all five original marker genes.

| Method | 1 | 2 | 3 | 4 |
| --- | --- | --- | --- | --- |
| ScType | 0.00 | 0.00 | 0.00 | 0.00 |
| scSorter | -23.46 | -23.98 | 0.00 | 0.84 |
| SCINA | -1.09 | -0.45 | -1.54 | 2.91 |

Overall, the marker genes used do impact our model’s performance. However, this impact is a result of decreased performance in the marker-based tools.

## G Comparison Between DeepLIFT and Differentially Expressed Genes

To compare highly attributed DeepLIFT genes and genes found through differential expression analysis, we used the Seurat FindAllMarkers() function. We used default parameters except for only.pos, which we set to true. This function uses a Wilcox based differential expression test to return differentially expressed genes for each cell type. Cell types assigned by our GCN model were used for this analysis.

Table 8 shows the top ten highly attributed DeepLIFT genes and top ten positively differentially expressed genes for all cell types in the testis. In Spermatogonia, seven of the top ten highly attributed DeepLIFT genes are also identified by differential expression. Similarly, six of the top ten highly attributed DeepLIFT genes for Spermatocytes are shared with differential expression analysis. Finally, in Spermatids, only five of the top ten highly attributed DeepLIFT genes are shared.

**Table 8.** Top 10 highly attributed DeepLIFT genes and top 10 positively differentially expressed (DE) genes for Spermatogonia (SPG), Spermatocytes (SPC), and Spermatids (SPD).

| Rank | SPG |  | SPC |  | SPD |  |
| --- | --- | --- | --- | --- | --- | --- |
|  | DeepLIFT | DE | DeepLIFT | DE | DeepLIFT | DE |
| 1 | mt-Rnr2 | Ncl | Ldhc | Rsph1 | Tnp1 | Smcp |
| 2 | Ncl | Hsp90ab1 | Ubb | Ldhc | Smcp | Tnp1 |
| 3 | Hsp90ab1 | Dazl | Fabp9 | Lyar | Tsga8 | Gm9999 |
| 4 | Dazl | Anp32b | Pabpc1 | Pabpc1 | Dbil5 | Fam229a |
| 5 | Prrc2c | Prrc2c | Meig1 | Morf4l1 | Gm9999 | Dbil5 |
| 6 | Tpr | Ssb | Calm2 | Cox8c | Acrv1 | Spata3 |
| 7 | Anp32b | Taf7l | Morf4l1 | Ccdc39 | Odf2 | DB30044l16Rik |
| 8 | mt-Nd1 | mt-Co1 | Calm1 | Calm1 | D830044l16Rik | Spata19 |
| 9 | Rpl4 | Rpl4 | Tuba3b | Cdc42ep3 | Ccdc136 | X1700029H14Rik |
| 10 | Rbm39 | Tpr | Rsph1 | Meig1 | mt-Rnr2 | X2610318N02Rik |

Table 9 shows the top ten highly attributed DeepLIFT genes and top ten positively differentially expressed genes for all cell types in the PBMC data set. For the top ten highly attributed genes in each cell type, seven, five, four, one, and three genes were shared with the top ten differentially expressed genes in B Cells, Monocytes, NK Cells, CD4 T Cells, and CD8 T Cells respectively. The low sharing between highly attributed and differentially expressed genes in T Cells likely stems from the low model accuracy identifying between CD4 and CD8 T Cells. We also found CD4 to be differentially expressed in CD4 T Cells after lowering the requirement for percentage of cells expressing the gene.

**Table 9.** Top 10 highly attributed DeepLIFT genes and top 10 positively differentially expressed (DE) genes for B Cells, Monocytes, Natural Killer Cells (NK Cells), CD4 T Cells (CD4), and CD8 T Cells (CD8). B Cells Monocytes NK Cells CD4 CD8

| Rank | B Cells |  | Monocytes |  | NK Cells |  | CD4 |  | CD8 |  |
| --- | --- | --- | --- | --- | --- | --- | --- | --- | --- | --- |
|  | DeepLIFT | DE | DeepLIFT | DE | DeepLIFT | DE | DeepLIFT | DE | DeepLIFT | DE |
| 1 | CD74 | HLA-DRA | FTH1 | S100A8 | GNLY | GNLY | LTB | IL32 | CD8B | CD8B |
| 2 | HLA-DRA | CD74 | TYROBP | S100A9 | NKG7 | NKG7 | FTH1 | AQP3 | CD3E | CD8A |
| 3 | CD79A | CD79A | S100A9 | CST3 | TYROBP | GZMB | CD3E | CD3E | CTSW | RP11-291B21-2 |
| 4 | CD79B | CD79B | S100A8 | LYZ | RPS21 | CLIC3 | MT-CO2 | LDHB | TPT1 | S100B |
| 5 | HLA-DRB1 | HLA-DPA1 | FTL | FCN1 | GZMB | FGFBP2 | RPL34 | MAL | EEF1A1 | CCL5 |
| 6 | LTB | HLA-DPB1 | CST3 | AIF1 | RPL38 | GZMA | S100A4 | CORO1B | HLA-C | GZMK |
| 7 | HLA-DPA1 | IGLL5 | LYZ | LST1 | HCST | CST7 | CD3D | JUNB | CCL5 | CARS |
| 8 | HLA-DPB1 | HLA-DRB1 | S100A4 | TYMP | FCER1G | FCER1G | RPL37 | CD27 | MALAT1 | CTSW |
| 9 | HLA-DRB5 | MS4A1 | HLA-DRB1 | TYROBP | FTL | KLRB1 | RPL7 | CD3D1 | CD3D | CPA5 |
| 10 | CD37 | HLA-DQA1 | UBC | CFD | RPS26 | PRF1 | UBC | IL7R | EEF1D | NELL2 |

## H Evaluation of Cell Type Similarities

To evaluate the difficulty of classifying our test data sets, we calculated the average distance between each cell type cluster for each data set. Figure 7 shows the results for this experiment in the form of heat maps. For each heat map, the scale goes from zero to the next highest integer above the highest average distance between any two cell types. This data shows that our simulated data sets consist of four cell types all with similar average distance from each other. One interesting aspect is that our Simulation 0.8 data set does not appear very different with this metric, despite being empirically easier to classify. In contrast to the simulated data sets, our testis and PBMC heat maps show varying similarities between cell type groups. For example. Spermatocytes and Spermatids have lower average distance than either of these cell types do with Spermatogonia. In the PBMC data set, CD4 and CD8 T Cells have lower average distance between each other than with other cell types.

**Fig. 7.**
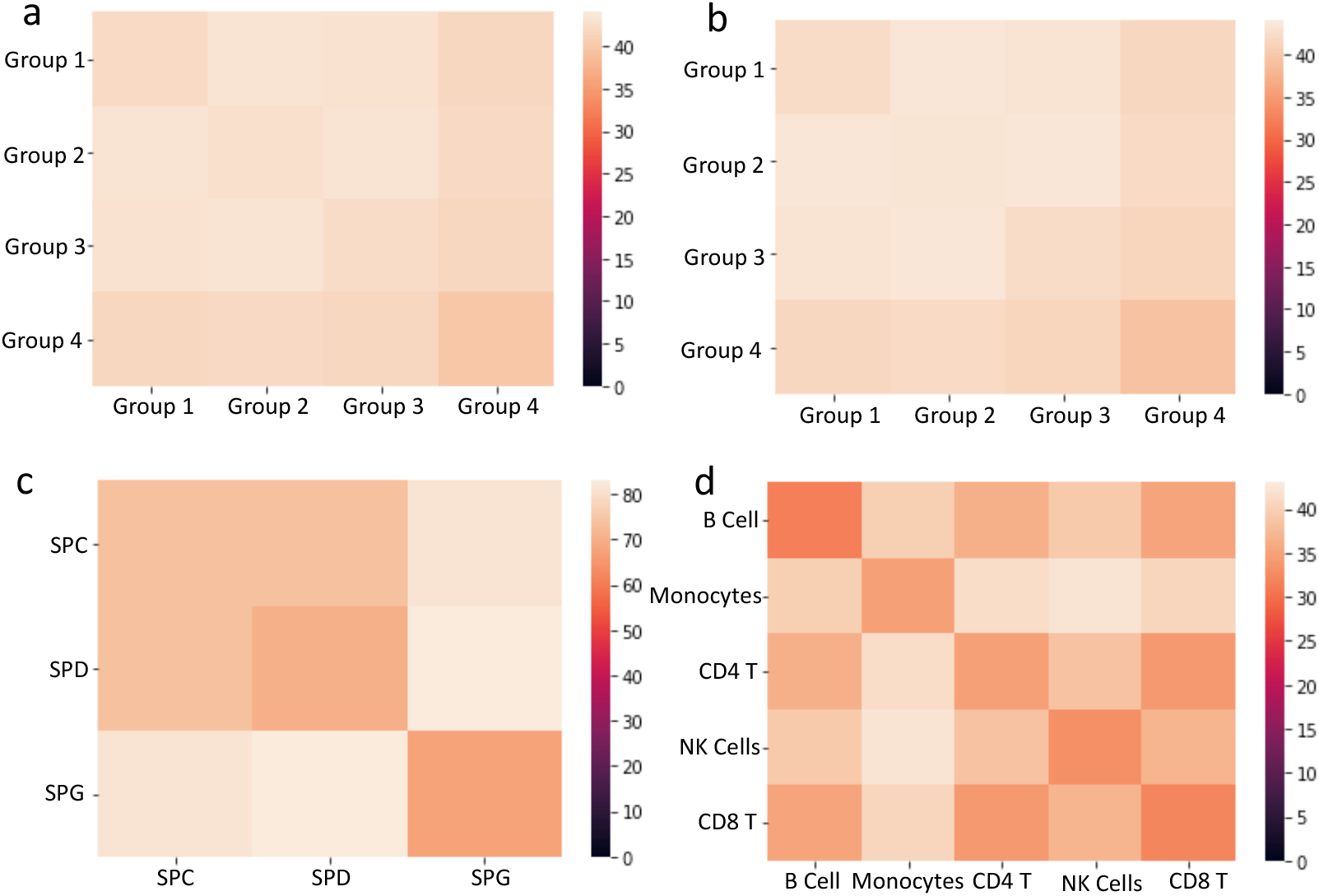
Heat maps showing the average distance between cell type clusters in each data set. a. Simulation 0.7 b. Simulation 0.8 c. Testis d. PBMC

In general, our simulated data sets show almost equal similarity between all cell types. In contrast, both real data sets show varying levels of similarity depending on which cell types are compared.

## I Marker Genes Used

The gene markers for the testis data set were as follows:

- Spermatogonia: Dazl, Sycp1
- Spermatocytes: Insl6, Piwil1, Pttg1, Spag6, Mllt10, Aurka
- Spermatids: Acrv1, Spaca1, Tsga8, Tssk6

The gene markers for the PBMC data set were as follows:

- B Cells: CD19, MS4A1, CD79A
- Monocytes: CD14, FCGR1A, CD68, S100A12
- Natural Killer Cells: NCAM1, FCGR3A
- CD4 T Cell: CD4, CD3D, CD3E, CD3G
- CD8 T Cell: CD8A, CD8B, CD3D, CD3E, CD3G

